# External validation and further exploration of fall prediction models based on questionnaires and daily-life trunk accelerometry

**DOI:** 10.1101/2022.09.26.509506

**Authors:** Yuge Zhang, Roel H.A. Weijer, Kimberley S. van Schooten, Sjoerd M. Bruijn, Mirjam Pijnappels

**Author notes:** Corresponding Author: Sjoerd M. Bruijn, Vrije Universiteit Amsterdam, De Boelelaan 1105, 1081 HV Amsterdam, Netherlands.

## Abstract

**Background:** Ambulatory measurements of trunk accelerations can provide valuable insight into the amount and quality of daily life activities. Such information has been used to create models that aim to identify individuals at high risk of falls. However, external validation of such prediction models is lacking, yet crucial for clinical implementation. We externally validated three previously described fall prediction models (van Schooten et al., 2015a).

**Methods:** Complete questionnaires and one week of trunk acceleration data were obtained in 263 community-dwelling people (mean age 71.8 years, 68.1% female). To validate models, we first used the coefficients and optimal cut-offs from original cohort, then recalibrated the original models, as well as optimized parameters based on our new cohort.

**Results:** Among all participants, 39.9% experienced falls during 6-month follow-up. All models showed poor precision (0.20-0.49), poor sensitivity (0.32-0.58), and good specificity (0.45-0.89). Calibration of the original models had limited effect on model performance. Using coefficients and cut-offs optimized on the external cohort also had limited benefit. Last, the odds ratios in our cohort were different from those in the original cohort and indicated that gait characteristics, except for index of harmonicity ML, were not statistically significantly associated with falls.

**Conclusions:** Prediction of fall risk in our cohort was not as effective as in the original cohort. Recalibration as well as optimized model parameters resulted in limited increase in accuracy. Fall prediction models are highly specific to the cohort studied. This highlights the need for large representative cohorts, preferably with an external validation cohort.

## 1. Introduction

Falls among older adults are a major cause of disability and death. Each year, one in three older adults aged 65 years or over experience at least one fall (Morrison et al. 2013; Salvà et al. 2004). Therefore, early identification of fall risk is necessary for timely intervention. Fall risk can be estimated by questionnaires, functional tests in a clinical setting or in a biomechanical laboratory, but all these methods have limited predictive value (Hamacher et al. 2011). Specifically, questionnaires have low reliability due to the subjective bias of the participants or testers (Furnham 1986; Malatesta et al. 2003; Winstein 1987). Clinical gait tests, such as Berg Balance scale, Timed Up and Go and POMA, cannot quantify fall risk well, especially in frail older populations (Barry et al. 2014; Boulgarides et al. 2003; Mancini & Horak 2010). Gait tests in a biomechanics laboratory do not reflect exposure to environmental and task-related hazards(Hausdorff et al. 2001; Maki 1997).

Inertial Measurement Units (IMU’s) are wearable motion sensors, which can be used in an everyday environment. Their validity and reliability have been shown in several studies. For instance, Felius and colleagues studied the test-retest reliability of 107 gait features assessed with IMUs (Felius et al. 2022). They found the relative minimal detectable change of these features ranged between 0.5 and 1.5 standard deviation. Several other studies compared IMUs to gold-standard gait-analysis technologies and suggested IMU-based gait metrics have sufficient accuracy and reliability to be used in clinical setting(Berner et al. 2020; Betteridge et al. 2021; Bezold et al. 2021; Zhong & Rau 2020).

Several studies have shown that daily-life gait quality characteristics, assessed with a single IMU, are able to discriminate fallers from non-fallers. Examples of such characteristics are local dynamic stability, variability of gait, frequency domain parameters, and gait smoothness (Bourke et al. 2010; Brodie et al. 2017; Rispens et al. 2015; Weiss et al. 2013). However, only a few research teams focus on developing fall risk prediction models based on daily-life activities in community environment (Brodie et al. 2017; Rispens et al. 2015; van Schooten et al. 2016; Weiss et al. 2013). Among these, three predictive models from van Schooten and colleagues (van Schooten et al. 2015a) had moderate to good predictive ability for falls happening in the next 6 months, and demonstrated that daily life accelerometry contributed substantially to the identification of individuals at risk of falls (areas under curve (AUC) of 0.68, 0.71 and 0.82 respectively). These models were logistic regression models based on a relatively large (n=169) data set of the FARAO cohort, a project concerning fall-risk assessment in older adults, with independent variables obtained either from questionnaires, from daily-life accelerometry, or from a combination of both.

However, to date, none of the fall-risk prediction models have been externally validated, so the repeatability and generalizability cannot be guaranteed (Nouredanesh et al. 2021), which hinders the implementation of a fall prediction model for community-dwelling older adults. Shany and colleagues (Shany et al. 2015) also pointed out that current studies of sensor-based fall risk estimation were overly optimistic due to small sample sizes, flawed modelling and validation methods, (e.g. inherent problems with overly cross-validation techniques, lack of external validation). In this study, we aimed to assess the external validity of the FARAO fall prediction models (van Schooten et al. 2015a) by using an external cohort of community-dwelling older adults. Moreover, as the ambulatory assessments specifically quantify the amount and quality of daily-life gait, we performed additional analyses that focused on walk-falls only. In addition, we calibrated the FARAO prediction models. Finally, we evaluated models based on the optimal model parameters of the new cohort and compared the odds ratio for individual variables between original and new cohorts.

## 2. Methods

### 2.1 Participants

The data in this study were available from the “Veilig in beweging blijven” (VIBE), which translates to “Safely remaining active” cohort. Participants were included if 1) they were 65 years of age or older, but not exceed 90 years 2) they had a Mini-Mental State Examination (MMSE) ≥ 19 (Folstein et al. 1975), 3) they were able to walk at least 20 meters, if necessary with a walking aid, without dizziness or perceiving chest pain or pressure, and 4) they were able to understand the Dutch language. The protocol was approved by the ethics committee of the faculty of Behavioural and Movement Sciences of the Vrije Universiteit Amsterdam (VCWE-2016-129). Among 441 persons who expressed interest, 283 were eligible to participate and agreed to a 12-month follow-up on falls. All participants provided signed informed consent before participation.

### 2.2 Questionnaire related to fall risk

Descriptive characteristics such as age, weight, height, the use of a walking aid and 6-month fall history were obtained from all participants. In addition, participants independently filled out validated questionnaires and tests on fall-risk factors, i.e. cognitive function (Mini Mental State Examination; MMSE), concern about falling (16-item fall efficacy scale; FES-I (Yardley et al. 2005)), LASA fall-risk profile (Longitudinal Aging Study Amsterdam) (Tromp et al. 2001) and Geriatric Depression Scale (GDS-15 (Yesavage et al. 1982)). The LASA fall-risk profile comprised questions concerning dizziness, independence in daily life, having pets, alcohol consumption, education duration and required the measurement of hand grip strength, which was quantified using a handgrip dynamometer (TKK 5401, Takei Scientific Instruments, Tokyo, Japan). The GDS-15 is an effective scale for depressive symptoms in older populations, where 5 negative items and 10 positive items indicate symptoms of depression (Greenberg 2012). Missing ≤20% of items in GDS-15 forms were imputed (Imai et al. 2014). Scores of 0-4 are considered normal, depending on age, education, and complaints; 5-8 indicate mild depression; 9-11 indicate moderate depression; and 12-15 indicate severe depression. Since the GDS-30 was used in the original model, we had to convert our GDS-15 scores to GDS-30 scores, according to a regression equation, GDS-30 scores = 1.95*GDS-15 scores + 1.57 (Zhang et al. 2023).

### 2.3 Assessment and analyses of daily-life gait characteristics

Conforming to the FARAO cohort study (van Schooten et al. 2015a), participants were instructed to wear a tri-axial accelerometer (DynaPort MoveMonitor, McRoberts, The Hague, the Netherlands) at home for 7 consecutive days, except during aquatic activities such as showering. The sensor was worn on the trunk at the level of L5 using an elastic belt, whose three axes of orientation corresponded to vertical (VT), anterior-posterior (AP) and medial-lateral (ML), ranged from −6g to +6g, with sampling frequency set to 100 Hz. We omitted the first 6 hours of the measurements from analysis, to discard any possible artefacts caused by transportation from the research team to the participants’ home. To have a valid measure of the amount of daily activity, the accelerometer had to be worn for more than twelve hours each day (van Schooten et al. 2015b). At the same time, the minimum valid days for daily physical activities are 2 days except for lying for which it is 3 days (van Schooten et al. 2015b), so only subjects who had accelerometer data with worn time ≥ 2 days were used. The identification of non-wearing, locomotion, sitting, lying, shuffling, and standing, were obtained from the manufacturer’s validated algorithms (Dijkstra et al. 2010). According to these episodes of daily activities, the duration of lying, walking, and the number of strides per day were calculated as outcome measures.

The data analyses were performed in Matlab R2019a (Mathworks, Natwick, MA), using the Gait Analysis toolbox (Human Movement Sciences 2022). Prior to estimation of gait characteristics, data from each locomotion episode was realigned with the vertical axis, which was estimated from the direction of the mean acceleration signal (Moe-Nilssen 1998), and anteroposterior and mediolateral directions, which were estimated by optimizing the left–right symmetry (Rispens et al. 2014). Locomotion episodes were then split into epochs of 10 seconds to avoid possible sample size-related bias in outcome measures (Bruijn et al. 2009). Median gait characteristics were calculated over epochs of each gait episode, identical to the data length requirements of characteristics determined, i.e., logarithmic divergence rate AP (Wolf et al. 1985), range VT, index of harmonicity ML (Lamoth et al. 2002) and sample entropy VT (Richman & Moorman 2000). The mean over all episodes in a week for these measures were calculated as outcome measure (van Schooten et al. 2016). Detailed description and formulas of sensor-based gait measures can be seen in the supplementary materials of (Felius et al. 2022).

### 2.4 Outcome: Falls and walk-falls

Although participants were monitored prospectively for falls over a 12-month period following the baseline assessment by means of a fall diary and monthly telephone contact, we only focused on the first 6 months of follow-up for consistency with the original models. However, to explore if extended follow-up periods resulted in better model performance, we also analyzed data obtained from the 12-month fall history and 12-month prospective fall recordings; these results are reported in the supplementary materials (see *Table S1-2* and *Fig. S1-2*). A fall was defined as an unintentional change in position resulting in coming to rest at a lower level or on the ground (Gibson 1987). Participants were classified as being a faller based on the number of prospectively monitored falls (≥1) they experienced over the 6-month period. Participants who did not record falls in their fall diary for the complete 6 months but experienced at least one fall within the recorded follow-up period were also classified as fallers.

When a fall happened, the activities and circumstances were also recorded, including walking, climbing the stair, riding a bicycle, showering, etc. Because the models were based on gait parameters, we also investigated how all models performed for people that fell specifically during walking. Participants were classified as walk-fallers if they had one record of a fall during walking.

### 2.5 Statistical analysis

Statistical analysis was performed in Matlab R2021a (MathWorks, Natick, MA) and R (R core team, Vienne). Unpaired t-tests were used to compare outcomes between fallers and non-fallers, and between walk-fallers and non-fallers, separately. The FARAO predictive models were all originally based on stepwise forward logistic regression analyses, with all continuous parameters transformed to z-scores as input. So, we did the same when we did our logistic regression. Model 1 only included 6-month history of falls and GDS-30, model 2 contained accelerometry-based daily life gait quantity and quality characteristics, and model 3 combined variables from both model 1 and model 2. Model performance was evaluated by area under the ROC curve (AUC), accuracy, precision, sensitivity, and specificity. Usually, AUC=0.5-0.7 means poor discriminative ability, 0.7-0.8 means moderate discriminative ability, 0.8-0.9 means good discriminative ability and 0.9-1.0 means excellent discriminative ability (Hosmer Jr et al. 2013).

To validate the FARAO predictive models on our VIBE cohort data, we first calculated model performance for our VIBE cohort when we used the mean/SDs of the FARAO cohort to calculate the z-scores for our data, and used the FARAO model coefficients, original cut-offs (0.279, 0.352 and 0.396). Either all falls or walking falls were used as outcome.

Next, these models were recalibrated (Van Calster et al. 2019) for possible model improvement. To this aim, the agreement between the predicted probability of the models and observed binary outcomes was evaluated using a logistic regression, in which the predicted probability was the independent variable and falling status the dependent variable, see equation 1

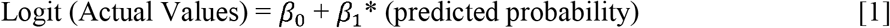

Where the slope (*β*_1_) indicates the level of overfitting (<1) or underfitting (>1) and the intercept (*β*_0_) suggests an general overestimation (<0) or underestimation (>0) (Van Calster et al. 2016). A slope of 1 and an intercept of 0 imply perfect model calibration (Van Calster et al. 2019). If these recalibrated models perform well, it suggests a simple scaling of the model would be enough to use the model in other populations.

Next, to see whether performance could be improved, considering potential differences in the sample measured, we fitted optimal logistic regression models for our VIBE dataset given the input variables as specified by the three FARAO models.

Finally, we performed unpaired t-tests on the means and SD of outcome measures of the VIBE data as compared to the published FARAO data, to see if potential differences in populations could explain performance. Lastly, univariate logistic regression was performed to identify the association between model variables and prospective falls, and these resulting odds ratio were compared with those from the FARAO cohort (van Schooten et al. 2015a).

## 3. Results

### 3.1 Descriptive Characteristics

283 participants filled in questionnaires and wore an accelerometer for one week at baseline, but 5 participants’ acceleration data could not be read. Among the remaining 278 participants, 15 participants were excluded due to missing more than 20% of the GDS items (n=3, 1 person missed 5 items and 2 persons missed 15 items), incomplete records within 6 months and no record of falls (n=3), accelerometer assessment duration of less than 2 days (n=3), ≤ 50 episodes of overall measurement time (n=6). Figure 1 presents a flowchart of the included fallers and non-fallers.

**Figure 1.**
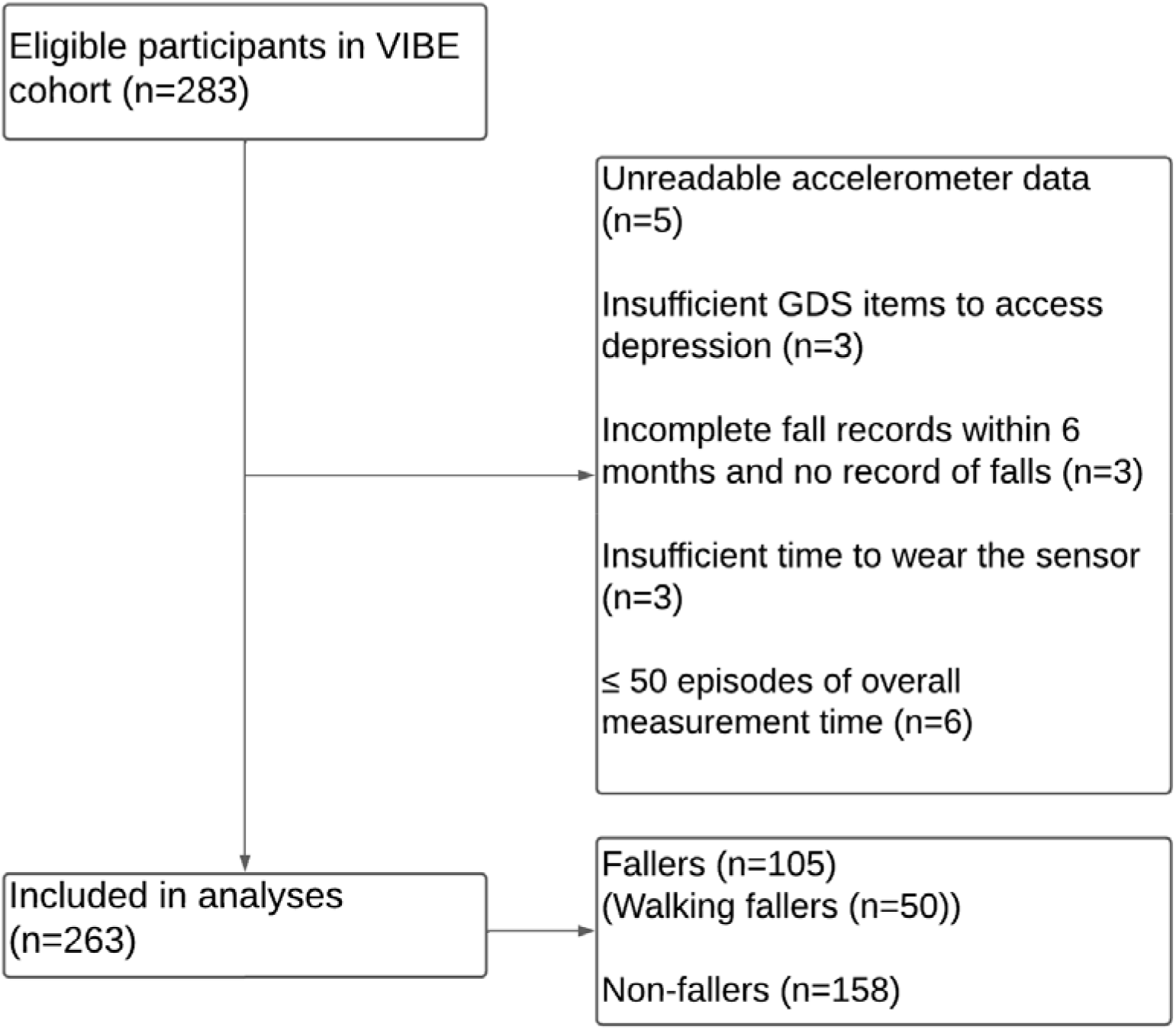
Flowchart of participants included for analyses. Fall records were considered incomplete when one or more months of fall records were missing.

Descriptive characteristics of the participants are summarized in table 1. Of the 263 participants, 39.9% (n=105) experienced one or more falls during follow-up, 19% (n=50) had fallen during walking. In addition, 37.6% of all participants, 43.8% of all fallers, 50% of walk-fallers and 33.5% of non-fallers had a fall history in the past 6 months. The average number of prospective and past falls, GDS scores, FES-I and LASA fall risk profile (score) were significantly different between fallers and non-fallers, and between walk-fallers and non-fallers. Furthermore, walk-fallers had significantly lower hand grip strength, and poorer independence in life, compared to non-fallers.

**Table 1.** Descriptive characteristics (Mean (SD)) of the VIBE participants

| Characteristics | All participants | All fallers | Walk-fallers | Non-fallers |
| --- | --- | --- | --- | --- |
| Number | 263 | 105 | 50 | 158 |
| Age (years) | 71.8 (5.7) | 71.9 (5.4) | 71.9 (6.2) | 71.8 (5.9) |
| Gender (%female) | 68.1 (179) | 68.6 (72) | 72 (36) | 67.7 (107) |
| Weight (kg) | 74.1 (13.5) | 72.6 (12.9) | 70.6 (12.4) | 75.1 (13.8) |
| Height (cm) | 169.1 (8.5) | 168.6 (9.1) | 166.4 (8.2) | 169.5 (8.1) |
| Number of falls in follow-up 6 months | 0.6 (1.02) | 1.6 (1.04)<sup>A</sup> | 2.0 (1.3)<sup>B</sup> | 0<sup>A, B</sup> |
| At least one fall in follow-up 6 months (%/n) | 39.9 (105) | 100 (105) | 100 (50) | 0 |
| Two or more falls in follow-up 6 months (%/n) | 14.5 (38) | 36.2 (38) | 56 (28) | 0 |
| Number of falls in past 6 months | 0.7 (1.4) | 1 (1.9)<sup>A</sup> | 1.4 (2.5)<sup>B</sup> | 0.5 (0.8)<sup>A, B</sup> |
| At least one fall in past 6 months (%/n) | 37.6 (99) | 43.8 (46) | 50 (25) | 33.5 (53) |
| Two or more falls in past 6 months (%/n) | 13.7 (36) | 20 (21) | 30 (15) | 9.5 (15) |
| Hand grip strength (kg) | 58.8 (16.6) | 57.5 (17.5) | 53.6 (17.5)<sup>b</sup> | 59.7 (15.9)<sup>b</sup> |
| Geriatric depression score 30, GDS (score) | 4.4 (4.3) | 5.5 (5.3)<sup>A</sup> | 6.5 (6.3)<sup>B</sup> | 3.7 (3.3)<sup>A, B</sup> |
| Fall efficacy scale, FES-I(score) | 20.5 (5.1) | 21.4 (6.5)<sup>a</sup> | 21.5 (6)<sup>b</sup> | 19.9 (3.9)<sup>a, b</sup> |
| Cognitive function (MMSE score) | 28.2 (1.6) | 28.3 (1.6) | 28.5 (1.7) | 28.2 (1.6) |
| LASA fall risk profile (score) | 3.4 (3.3) | 4.4 (3.8)<sup>A</sup> | 5.4 (4.5)<sup>B</sup> | 2.8 (2.8)<sup>A, B</sup> |
| Inability of using transport(%/n)<sup>1</sup> | 0.4 (1) | 1 (1) | 0 (0) | 0 (0) |
| Living independently (%/n) | 99.2 (261) | 98.1 (103) | 96 (48)<sup>b</sup> | 100 (158)<sup>b</sup> |
| Use of a walking aid (%/n) | 6.8 (18) | 8.6 (9) | 10 (5) | 5.7 (9) |

**Note:** <sup>1</sup>Either public or own transport. <sup>A</sup> significant difference between all fallers and non-fallers with p<0.01; <sup>a</sup> significant difference between all fallers and non-fallers with p<0.05; <sup>B</sup> significant difference between walking fallers and non-fallers with p<0.01; <sup>b</sup> significant difference between walking fallers and non-fallers with p<0.05.

Figure 2(A) shows the main activities when a fall happened, including walking (39%), riding a bicycle (13%), movement transfer (12%), going up or down the stairs (12%), doing sports (6%), standing, and sitting (in static positions) (4%), and other activities (15%). Among these, hiking was classified as a sport in this study. Working in the garden, getting dressed, etc. were classified as “Other”. Furthermore, main causes of falls during walking were shown in figure 2(B), which were trip/stumble (34%), balance loss (31%), slip (17%), physiological discomfort (8%), being distracted (2%) and other (8%). Trip/stumble often happened on uneven roads or roads with obstacles, and balance loss was more likely to occur when there was sudden disturbance, opposite walking height to the expected. Falling due to slips usually occurred in the bathroom or on a slippery floor. In addition, physiological discomfort included ankle sprain, sudden dizziness, drops in blood pressure, eye injury and weakened muscle strength, etc. Falling caused by distraction included chatting, looking at map, or doing other tasks when walking. Lastly, some unknown, poorly classified reasons were divided into “Other”.

**Figure 2.**
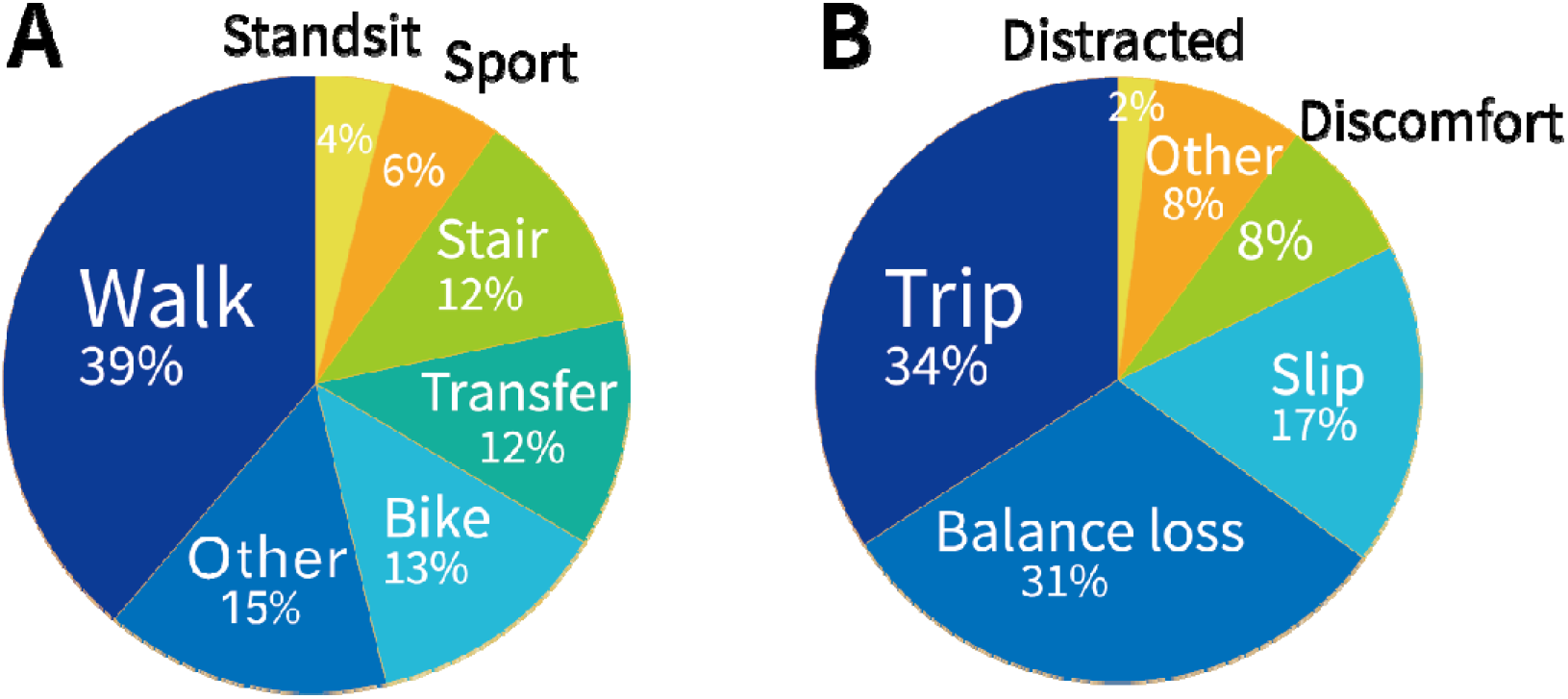
Main activities of all falls (A) and causes of walk-falls (B)

### 3.2 Validation of predicting all falls and walk-falls based on existing models

By using optimal cut-offs (0.279, 0.352 and 0.396), coefficients and mean (SD) (for z-score) from the original cohort, we obtained model performance of the three prediction models based on only questionnaires, only accelerometry, and combined questionnaires and accelerometry. Table 2a shows the results of each prediction model, with either all falls or walk-falls as outcome (Model performance for 12-months follow-up on FARAO cohort can be seen in *Table S2a*). Model 1, with only questionnaires, performed best for predicting all falls and walk-falls, with AUC (area under the curve) of 0.62 and 0.68, respectively. AUC of model 2 in all-fall and walk-fall groups was 0.50 and 0.49, respectively, and 0.53 and 0.59 in model 3. At the same time, the precision and sensitivity in both groups were low, which meant the ability of predicting a fall was low for all models.

**Table 2.** Model performance of fallers in follow-up 6 months for a) models based on the FARAO cohort, b) the calibrated FARAO model performance, c) models based on VIBE cohort

| a) model based on FARAO cohort | All falls |  |  | Walk-falls |  |  |
| --- | --- | --- | --- | --- | --- | --- |
|  | Model 1 | Model 2 | Model 3 | Model 1 | Model 2 | Model 3 |
| Accuracy | 0.58 | 0.50 | 0.49 | 0.63 | 0.53 | 0.75 |
| Precision | 0.48 | 0.37 | 0.40 | 0.34 | 0.20 | 0.49 |
| Sensitivity | 0.50 | 0.36 | 0.55 | 0.58 | 0.32 | 0.34 |
| Specificity | 0.64 | 0.59 | 0.45 | 0.64 | 0.59 | 0.89 |
| AUC | 0.62 | 0.50 | 0.53 | 0.68 | 0.49 | 0.59 |
| b) calibrated FARAO model performance | All falls |  |  | Walk-falls |  |  |
|  | Calibrated Model1 | Calibrated Model2 | Calibrated Model3 | Calibrated Model1 | Calibrated Model2 | Calibrated Model3 |
| Optimal cut-offs | 0.36 | 0.39 | 0.45 | 0.23 | 0.24 | 0.30 |
| Accuracy | 0.59 | 0.48 | 0.63 | 0.71 | 0.57 | 0.75 |
| Precision | 0.49 | 0.42 | 0.59 | 0.42 | 0.27 | 0.47 |
| Sensitivity | 0.53 | 0.86 | 0.28 | 0.50 | 0.48 | 0.36 |
| Specificity | 0.63 | 0.22 | 0.87 | 0.78 | 0.59 | 0.87 |
| AUC | 0.62 | 0.52 | 0.58 | 0.68 | 0.51 | 0.62 |
| c) model based on VIBE cohort | All falls |  |  | Walk-falls |  |  |
|  | Model 1 | Model 2 | Model 3 | Model 1 | Model 2 | Model 3 |
| Optimal cut-offs | 0.38 | 0.40 | 0.37 | 0.20 | 0.23 | 0.20 |
| Accuracy | 0.62 | 0.53 | 0.57 | 0.66 | 0.54 | 0.6 |
| Precision | 0.53 | 0.43 | 0.47 | 0.38 | 0.27 | 0.33 |
| Sensitivity | 0.56 | 0.53 | 0.58 | 0.66 | 0.54 | 0.62 |
| Specificity | 0.66 | 0.53 | 0.56 | 0.66 | 0.54 | 0.59 |
| AUC | 0.63 | 0.53 | 0.65 | 0.69 | 0.56 | 0.73 |

### 3.3 Model recalibration

In our VIBE cohort, the actual fall incidence and walk-fall incidence in 6 months was 0.40 and 0.19. The recalibrated fall rate was the same as actual fall rate, but the recalibrated walk-fall rate was 0.24 (see *Table S3*). Calibration intercepts of model 1 and model 3 in the group with all-fallers were significantly negative (−1.06 and −0.83, respectively), as well as intercepts of all models with walk-fallers (−2.17, −1.14 and −1.82, respectively). The slopes of model 1 and model 3 were significantly >1 in both groups (2.22 and 1.40, 3.24 and 2.08, respectively), indicating underfitting. The performance of the calibrated models is shown in table 2b. The AUC of all calibrated models ranged from 0.51-0.68.

### 3.4 Validation of predicting all falls and walk-falls based on the external cohort

Model performances did not improve as compared to the original FARAO model when the models were fitted again (with the same variables as in the original FARAO model), shown in table 2c (see ROC curves in *Fig. S3*) (12-months follow-up model performance and ROC curves based on VIBE cohort can be seen in *Table S2b* and *Fig. S2*, respectively). Specificity and sensitivity were still poor for these models (ranging from 0.59-0.66).

### 3.5 Comparison of cohorts and individual odds ratios

To better understand differences in model performance between the original models and models fitted on our external dataset, the comparison of descriptive statistics and odds ratio of measures for the our external VIBE cohort and original FARAO cohort (van Schooten et al. 2015a) are presented in table 3. There were small but significant differences in age, and height, with the VIBE cohort being slightly younger and shorter. Prevalence of fall history prior to 6 months in VIBE cohort was 37.6%, which was slightly higher than 35.5% in the FARAO cohort. Cognitive function (MMSE score) of VIBE cohort was 28.2, slightly higher than 27.7 in FARAO cohort. LASA fall-risk profile score, percent of inability to use public transportation and use of walking aid in VIBE cohort were lower than those in FARAO cohort. In agreement with this, the participants of the VIBE cohort had a higher degree of living independence. Number of strides walked, and walking duration were significantly higher in the VIBE cohort than in the FARAO cohort. However, subjects in the VIBE cohort walked with lower walking speed (0.6 m/s for VIBE, vs 0.9 m/s for FARAO). The duration of lying was 0.9 hours less than FARAO cohort. All gait characteristics differed significantly between the VIBE and FARAO cohorts.

**Table 3.**
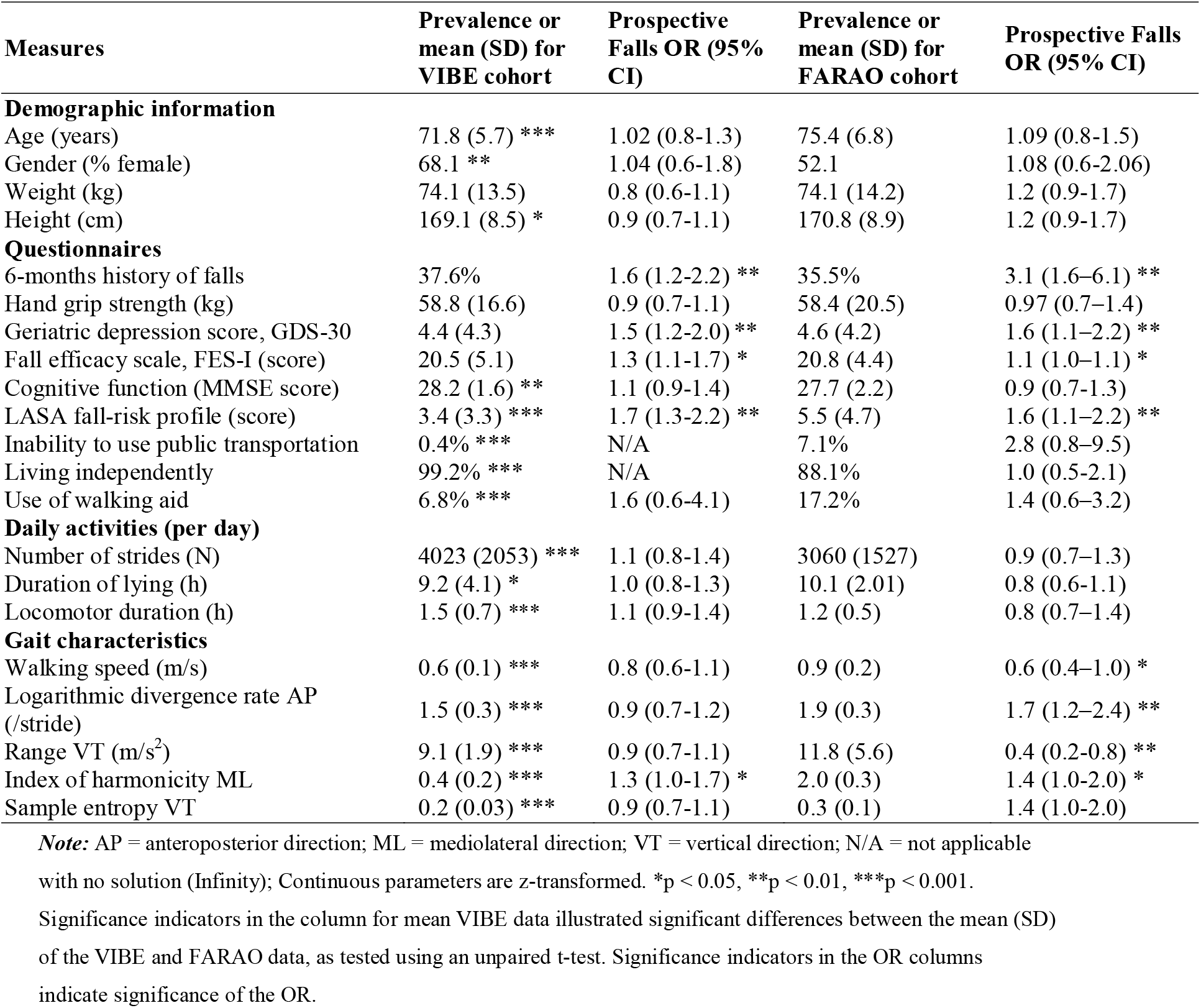
Mean (SD) and odds ratios (OR) for VIBE cohort (n=263) and FARAO cohort (n=169) (van Schooten et al. 2015a)

| Measures | Prevalence or mean (SD) for VIBE cohort | Prospective Falls OR (95% CI) | Prevalence or mean (SD) for FARAO cohort | Prospective Falls OR (95% CI) |
| --- | --- | --- | --- | --- |
| **Demographic information** |  |  |  |  |
| Age (years) | 71.8 (5.7) *** | 1.02 (0.8-1.3) | 75.4 (6.8) | 1.09 (0.8-1.5) |
| Gender (% female) | 68.1 ** | 1.04 (0.6-1.8) | 52.1 | 1.08 (0.6-2.06) |
| Weight (kg) | 74.1 (13.5) | 0.8 (0.6-1.1) | 74.1 (14.2) | 1.2 (0.9-1.7) |
| Height (cm) | 169.1 (8.5) * | 0.9 (0.7-1.1) | 170.8 (8.9) | 1.2 (0.9-1.7) |
| **Questionnaires** |  |  |  |  |
| 6-months history of falls | 37.6% | 1.6 (1.2-2.2) ** | 35.5% | 3.1 (1.6-6.1) ** |
| Hand grip strength (kg) | 58.8 (16.6) | 0.9 (0.7-1.1) | 58.4 (20.5) | 0.97 (0.7-1.4) |
| Geriatric depression score, GDS-30 | 4.4 (4.3) | 1.5 (1.2-2.0) ** | 4.6 (4.2) | 1.6 (1.1-2.2) ** |
| Fall efficacy scale, FES-I (score) | 20.5 (5.1) | 1.3 (1.1-1.7) * | 20.8 (4.4) | 1.1 (1.0-1.1) * |
| Cognitive function (MMSE score) | 28.2 (1.6) ** | 1.1 (0.9-1.4) | 27.7 (2.2) | 0.9 (0.7-1.3) |
| LASA fall-risk profile (score) | 3.4 (3.3) *** | 1.7 (1.3-2.2) ** | 5.5 (4.7) | 1.6 (1.1-2.2) ** |
| Inability to use public transportation | 0.4% *** | N/A | 7.1% | 2.8 (0.8-9.5) |
| Living independently | 99.2% *** | N/A | 88.1% | 1.0 (0.5-2.1) |
| Use of walking aid | 6.8% *** | 1.6 (0.6-4.1) | 17.2% | 1.4 (0.6-3.2) |
| **Daily activities (per day)** |  |  |  |  |
| Number of strides (N) | 4023 (2053) *** | 1.1 (0.8-1.4) | 3060 (1527) | 0.9 (0.7-1.3) |
| Duration of lying (h) | 9.2 (4.1) * | 1.0 (0.8-1.3) | 10.1 (2.01) | 0.8 (0.6-1.1) |
| Locomotor duration (h) | 1.5 (0.7) *** | 1.1 (0.9-1.4) | 1.2 (0.5) | 0.8 (0.7-1.4) |
| **Gait characteristics** |  |  |  |  |
| Walking speed (m/s) | 0.6 (0.1) *** | 0.8 (0.6-1.1) | 0.9 (0.2) | 0.6 (0.4-1.0) * |
| Logarithmic divergence rate AP (/stride) | 1.5 (0.3) *** | 0.9 (0.7-1.2) | 1.9 (0.3) | 1.7 (1.2-2.4) ** |
| Range VT (m/s<sup>2</sup>) | 9.1 (1.9) *** | 0.9 (0.7-1.1) | 11.8 (5.6) | 0.4 (0.2-0.8) ** |
| Index of harmonicity ML | 0.4 (0.2) *** | 1.3 (1.0-1.7) * | 2.0 (0.3) | 1.4 (1.0-2.0) * |
| Sample entropy VT | 0.2 (0.03) *** | 0.9 (0.7-1.1) | 0.3 (0.1) | 1.4 (1.0-2.0) |

**Note:** AP = anteroposterior direction; ML = mediolateral direction; VT = vertical direction; N/A = not applicable with no solution (Infinity); Continuous parameters are z-transformed. \*p < 0.05, \*\*p < 0.01, \*\*\*p < 0.001. Significance indicators in the column for mean VIBE data illustrated significant differences between the mean (SD) of the VIBE and FARAO data, as tested using an unpaired t-test. Significance indicators in the OR columns indicate significance of the OR.

For both the VIBE and the FARO cohorts, significant odds ratios for questionnaire metrics 6-months history of falls, GDS-30, FES-I, and LASA profile score were found. For the VIBE cohort, the only gait measure that was significantly associated with falls was the index of harmonicity ML. However, the confidence interval of the odds ratios for the other gait parameters overlapped with those in FARAO cohort, except for the logarithmic divergence rate AP.

## 4. Discussion

We validated the predictive ability of three fall prediction models based on questionnaires, accelerometry of daily life gait, and both, on the data from the VIBE cohort of community-dwelling older adults. We found that the external validity of the models developed based on the FARAO cohort was limited for predicting all falls as well as when focusing on walk-falls. Subsequently, we explored why this may have been so.

When comparing the population characteristics between the two cohorts, we found that participants in the VIBE cohort were slightly younger, more often female, a little shorter, had a higher level of cognition, had less fall risk, lived more independently, use public transport more often and used less walking aids. Participant’s in both studies had a normal (i.e., low) degree of depressive symptoms (Greenberg 2012). Both cohorts had the same definition of a faller in past that the number of falls in follow-up 6 months was ≥1. The prevalence of falls in our study was higher than in the original study (39.9% and 34.9%, respectively), with 14.5% participants having two or more falls in the follow-up 6 months compared to 16.5% in the FARAO cohort. Studies suggested that only one fall may be an ambiguous fall, with no apparent cause (Nouredanesh et al. 2021), and people who were ‘once only fallers’ were characteristically more closely related to non-fallers than ‘recurrent fallers’(Lord et al. 1994) or injury-prone fallers. Hence, the fall rate of the FARAO cohort was lower than VIBE cohort if based on one fall, but higher than the VIBE cohort if based on multiple falls.

The definition and record methods of falls were the same in both cohorts, including fall calendars and monthly phone calls. However, this does not exclude that there was a slight difference in interpreting the fall during the phone calls that may have resulted in more reported falls on the VIBE cohort. Causes of falls were divided into 7 categories and 39%of falls occurred during walking, similar to other studies (Niino et al. 2000; Peel 2011; Talbot et al. 2005). However, it was not known if falls in the FARAO cohort had a similar distribution of causes. Hence the different proportion of falls during walking may have affected the results, but the models also did not work well for the walking fallers. In post-hoc analysis, results did not change based on the 12-month follow-up data available from the VIBE cohort (refer to *Table S2*). In conclusion, there may have been differences in the classification of fallers that influenced the accuracy of the models. However, since such differences are most likely due to interpretation of the fall definition by the people who performed the monthly phone calls, we are unable to identify such potential differences to test if these affected the results.

Limited model performance in both groups was obtained after recalibration of the model from the FARAO cohort. Calibrated slopes were significantly larger than 1 and intercepts were significantly negative. This indicates underfitting and overestimation of original models. Some studies recommend use dataset with more than 100 events and 100 non-events to prevent the potential underfitting of models (Van Calster et al. 2016), and it is clear that the original FARAO cohort, with 169 subjects, does not fulfill this criterium.

When we explored the optimal coefficients and optimal cut-offs for the VIBE cohort, we found that model 3 performed best for both all falls and walk-falls. The AUC of all models improved a little, but the precision and sensitivity remained moderate. More importantly, the model which combined the gait quality characteristics and questionnaires was better than either one alone – reinforcing the conclusion of van Schooten et al.’s study that daily-life accelerometry has value for the identification of people at risk for falls. Thus, while the same variable may yield a reasonable fall prediction model, it appears that coefficients and cut-offs may be rather specific to a given population sample (Punt et al. 2016).

To further study the predictors in the models, odds ratios of individual variables were compared with the initial FARAO cohort. Studies have shown that fall history, depressive symptoms, LASA fall-risk profile, fear of falling, and poor executive function (as assessed by TMT-A) were associated with prospective falls (Holtzer et al. 2007; Pluijm et al. 2006; Tromp et al. 2001). Significant odds ratios of these measures in the original study confirmed this (van Schooten et al. 2015a). However, of the gait quality characteristics, only index of harmonicity ML had significant odds ratios in VIBE cohort, while other gait quality measures have been shown to be related to future falls in FARAO cohort. Specifically, lower harmonic ratio, reflecting lower gait symmetry, was shown to be related to falls by (Doi et al. 2013); higher logarithmic rate of divergence, reflecting lower local stability, was associated with future falls, according to previous findings (Lockhart & Liu 2008; Rispens et al. 2015; Toebes et al. 2012). This was most likely due to the differences in the VIBE cohort, who were younger, and more active than participants in the FARAO cohort. Still, these findings suggest that even though we could not successfully use the FARAO fall prediction models for an external data set, there may be merit in the use of daily-life IMU data.

Our study has some limitations. First, we included elderly subjects who did or not use walking aids, but it is uncertain whether the use of a walking aid influenced the extracted gait characteristics, and their relationship with fall risks, which is also a limitation in the original FARAO models. Second, we were unable to pinpoint the source of some large differences in gait parameters between our cohort. For instance, we found a complete lack of overlap for the index of harmonicity ML in the mean values and its standard deviation (0.4(0.2) and 2.0 (0.3)). However, this also highlights the inherent difficulty with creating widely applicable fall-prediction models; even though our inclusion criteria were largely similar, and we used similar recruitment procedures, our results were substantially different from previous studies. Perhaps, even larger studies (or pooling of datasets) can prevent these shortcomings.

## 5. Conclusion

In conclusion, prediction of fall risk in our VIBE cohort based on the FARAO models was not as effective as models in FARAO cohort (van Schooten et al. 2015a). Moreover, recalibration of the models suggested that the original models are underfitting and have limited external validity. Despite several earlier reports indicating the importance of daily life gait measures for predicting falls, such measures alone were of limited value in our cohort. Moreover, the fact that optimal coefficients and cut-off were different for our cohort, as well as the fact that we found almost no significant ORs for the gait parameters included in the model, suggest that this kind of fall prediction models are highly specific to the cohort studied. Nonetheless, even with these optimal coefficients and cut-offs, model performance was still poor, which highlights the need for large representative cohorts, preferably with an external validation cohort.

## Author contributions

Conceptualization, MP and SMB; methodology, RW, SMB, KS and MP; formal analysis, YZ, SMB, MP, RW and KS; resources, MP and SMB; data collection, RW; data processing, YZ; writing-review and editing, YZ, RW, SMB, KS and MP; visualization, YZ; supervision, SMB and MP; project administration, MP. All authors have read and agreed to the published version of the manuscript.

