## supplementary figure 3, supplementary table 2b and figure 2, supplementary tables 1-2 and figures 1-2, supplementary table 2a, supplementary table 3 for "External validation and further exploration of fall prediction models based on questionnaires and daily-life trunk accelerometry"

### Supplementary materials

**Supplementary table 1. Descriptive characteristics (Mean (SD)) of the participants with prospective 12-month falls**

| **Characteristics** | All participants | All fallers | Walk-fallers | Non-fallers |
| --- | --- | --- | --- | --- |
| Number | 250 | 149 | 84 | 101 |
| Age (years) | 71.7 (5.7) | 71.5 (5.5) | 71.6 (5.9) | 71.9 (6) |
| Gender (%female) | 67.6 (169) | 69.8 (104) | 77.4 (65) | 64.4 (65) |
| Weight (kg) | 73.8 (13.2) | 73.7 (13.2) | 71.6 (12.7) | 74 (13.4) |
| Height (cm) | 169.2 (8.6) | 169 (8.7) | 167.7 (8.1) | 169.4 (8.4) |
| Number of Falls in follow-up 12 months | 1.3 (1.8) | 2.2 (2.0) | 2.8 (2.5) | 0 |
| At least one fall in follow-up 12 months (%/n) | 59.6% (149) | 100 (149) | 100 (84) | 0 (0) |
| Two or more falls in follow-up 12 months (%/n) | 31.6 (79) | 53.0 (79) | 69.1 (58) | 0 (0) |
| Number of falls in past 6 months | 0.7 (1.4) | 1 (1.9) ^A^ | 1.4 (2.5) ^B^ | 0.5 (0.8) ^A, B^ |
| At least one fall in past 6 months (%/n) | 37.6 (99) | 43.8 (46) | 50 (25) | 33.5 (53) |
| Two or more falls in past 6 months (%/n) | 13.7 (36) | 20 (21) | 30 (15) | 9.5 (15) |
| Hand grip strength (kg) | 59.1 (16.7) | 58.7 (17.1) | 55.5 (16) | 59.8 (16.2) |
| Geriatric depression score, GDS-30 | 4.4 (4.3) | 5.1 (4.9) ^A^ | 5.8 (5.6) ^B^ | 3.3 (3.0) ^A, B^ |
| Fall efficacy scale, FES-I(score) | 20.5 (5.2) | 20.8 (5.7) | 20.9 (5.3) | 20 (4.3) |
| LASA fall risk profile (score) | 3.5 (3.3) | 4.2 (3.6) ^A^ | 5.1 (4) ^B^ | 2.5 (2.6) ^A, B^ |
| Cognitive function (MMSE score) | 28.3 (1.6) | 28.4 (1.6) | 28.5 (1.6) | 28.1 (1.5) |
| Inability of using transport(%/n)^1^ | 0.4 (1) | 0.7 (1) | 0 (0) | 0 (0) |
| Living independently (%/n) | 99.2 (248) | 98.7 (147) | 97.6 (82) | 100 (101) |
| Use of a walking aid (%/n) | 6.4 (16) | 8.1 (12) | 8.3 (7) | 4 (4) |

***Note:*** ^1^Either public or own transport. A: significant difference between all fallers and non-fallers with p<0.01; a: significant difference between all fallers and non-fallers with p<0.05; B: significant difference between walking fallers and non-fallers with p<0.01. b: significant difference between walking fallers and non-fallers with p<0.05.

**Supplementary table 2. Model performance of fallers in follow-up 12 months for a) models based on original FARAO cohort, b) models based on VIBE cohort**

| **a) model based on FARAO cohort** | **All falls** | | | | **Walk-falls** | | |
| --- | --- | --- | --- | --- | --- | --- | --- |
|  | **Model1** | **Model2** | | **Model3** | **Model1** | **Model2** | **Model3** |
| Accuracy | 0.57 | 0.45 | 0.51 | | 0.64 | 0.45 | 0.48 |
| Precision | 0.70 | 0.56 | 0.60 | | 0.60 | 0.37 | 0.44 |
| Sensitivity | 0.50 | 0.36 | 0.55 | | 0.58 | 0.30 | 0.51 |
| Specificity | 0.68 | 0.58 | 0.46 | | 0.68 | 0.58 | 0.46 |
| AUC | 0.65 | 0.47 | 0.54 | | 0.71 | 0.44 | 0.51 |
| **b) model based on VIBE cohort** | **All falls** | | | | **Walk-falls** | | |
|  | **Model1** | **Model2** | | **Model3** | **Model1** | **Model2** | **Model3** |
| Optimal cut-offs | 0.58 | 0.6 | | 0.56 | 0.2 | 0.23 | 0.2 |
| Accuracy | 0.61 | 0.53 | | 0.63 | 0.66 | 0.54 | 0.6 |
| Precision | 0.69 | 0.62 | | 0.72 | 0.38 | 0.27 | 0.33 |
| Sensitivity | 0.63 | 0.52 | | 0.62 | 0.66 | 0.54 | 0.62 |
| Specificity | 0.58 | 0.53 | | 0.63 | 0.66 | 0.54 | 0.59 |
| AUC | 0.66 | 0.53 | | 0.67 | 0.69 | 0.56 | 0.73 |

**Supplementary table 3. Calibration slope and intercept for follow-up 6-month fallers**

| **Coefficient** | **All falls** | | | **Walk-falls** | | |
| --- | --- | --- | --- | --- | --- | --- |
|  | **Model1** | **Model2** | **Model3** | **Model1** | **Model2** | **Model3** |
| Fall incidence | 0.40 | | | 0.19 | | |
| Predicted fall incidence | 0.40 | | | 0.24 | | |
| intercept | -1.06 *** | -0.56 | -0.83 *** | -2.17 *** | -1.14 * | -1.82 *** |
| slope | 2.22 ** | 0.51 | 1.40 ** | 3.24 *** | - 0.03 | 2.08*** |

***Note:*** *p < 0.05, **p < 0.01, ***p < 0.001.
